# DANEELpath: Open-source Digital analysis tools for histopathological research. Applications in NEuroblastoma modELs

**DOI:** 10.1101/2025.03.31.646267

**Authors:** Isaac Vieco-Martí, Amparo López-Carrasco, Samuel Navarro, Sofia Granados Aparici, Rosa Noguera

## Abstract

Despite the considerable expansion of bioimage analysis as a subfield of biomedical sciences, there is an ongoing need for comprehensive image analysis pipelines to address specific biological inquiries. In the tumor microenvironment, the extracellular matrix (ECM) plays a pivotal role in cancer progression, promoting tumor cell adaptability, intratumor heterogeneity, and therapeutic resistance. In neuroblastoma (NB), the ECM glycoprotein vitronectin (VN) has been associated with more aggressive tumors. Three-dimensional (3D) hydrogels are an emerging biomimetic tool with significant potential for studying the role of ECM elements and testing new mechano-drugs such as cilengitide (CLG), a potential therapeutic agent to treat high-risk (HR) NB due to its ability to inhibit VN activity in cells. To gain a more detailed understanding of the effects of VN and CLG in 3D-grown NB cells, we developed DANEELpath, an open-source image analysis toolkit. DANEELpath integrates deep learning techniques, specific segmentation of individual and cluster cells through mathematical morphology pipelines, and extraction of spatial features within whole-slide images. Thanks to its versatility, DANEELpath is adaptable to address different biological questions and has significant potential for use in a variety of research fields and model systems, which could help advance biomedical discovery.

## INTRODUCTION

Bioimage analysis is experiencing significant growth within the biomedical sciences field as it provides a means of conducting objective and reproducible quantification of histological images (1). This has led to an increasing demand for specific image analysis pipelines and user-friendly tools for detailed analysis of specific biological questions (2,3).

The development of detailed protocols for digital image analysis of histological slides and new deep learning tools such as StarDist (4), CellPose (5) and InstanSeg (arXiv:2408.15954) has become essential to study the complex behavior of tumor cells in a spatio-temporal context related to the tumor microenvironment (TME). The TME plays a pivotal role in cancer progression, promoting tumor cell adaptability, intratumor heterogeneity and therapeutic response (6). Within the TME, the Extracellular Matrix (ECM) forms a complex network of fibrillar proteins, glycoproteins, and proteoglycans in cells to connect the cytoskeleton (7,8). These cell connections to the ECM (through receptors such as integrins) trigger mechanotransduction processes that affect gene expression, and cell survival, progression, and migration (9,10).

Neuroblastoma (NB) is the most prevalent extracranial solid tumor of childhood (11), with a five-year survival rate for high-risk NB (HR-NB) patients of under 50% (12). Among the biomarkers of this disease is Vitronectin (VN), a cell-ECM anchorage glycoprotein that interacts strongly with integrins αvβ3/αvβ5, uPAR, PAI-1 and collagens (13,14). Presence of VN in biopsy samples has been linked to poor prognosis, and a recent study has also reported elevated VN levels in the plasma of NB patients (15,16). In terms of study methods, the advent of three-dimensional (3D) cell culture has driven the development of systems that more accurately mimic the architecture and dynamics of tumors. SK-N-BE (2) and SH-SY5Y NB cell lines have been efficiently grown in polyethylene glycol (PEG) and gelatin/methacrylate/alginate (GelMaAlgM)-based hydrogels (17,18). Recently, gelatin-tyramine with silk fibroin models (GTA-sf) have also been shown to promote growth in NB and Ewing sarcoma cell lines (19). GTA-sf models facilitate VN incorporation into the matrix, avoiding the need for complex chemical processes. To prevent interactions between the ECM and/or cells and VN in NB cells, researchers have used cilengitide (EMD 121974, CLG), a mechano-drug based on cyclo(-RGDfV-), a cyclic peptide which is selective for ανβ3 and ανβ5 integrins in HR-NB cells (20,21).

A recent study using digital holographic imaging with GTA-sf 3D models over a 10-day culture period revealed different growth dynamics over time and a differential CLG response in cell lines with different genetic backgrounds (22). While digital holography proved invaluable for studying growth dynamics in real time, the underlying mechanisms driving these behaviors could not be identified with this method. Formalin-Fixed Paraffin-Embedded (FFPE) GTA-sf hydrogel sections provide a valuable opportunity to explore the effects of CLG as regards the spatial distribution of cells within matrices, as well as the expression of molecules such as VN through immunohistochemical assays. By digitizing samples as whole slide images (WSI) we can use digital image analysis tools to obtain morphological features.

Previous studies using hydrogels for NB cells have demonstrated a capacity of cells to proliferate from a few initial cells to hundreds within a cell cluster formation (17,18). Furthermore, WSI presents constraints in terms of extracting staining features for specific markers and spatial information pertaining to inner cells and cell clusters, often due to the lack of a standardized protocol for histopathological digital analysis. These limitations preclude increasing sample sizes or testing a greater number of mechano-drugs in tumor models. To address this issue, we developed DANEELpath, an open-source toolkit for histopathological digital analysis for NB models.

This study outlines the functionality of DANEELpath tools, demonstrating their role in facilitating the process of cluster segmentation and spatial data acquisition. Furthermore, we provide a pipeline for extracting morphological features of pericluster structures. DANEELpath tools are designed to function with QuPath, an open-source software widely used in digital pathology (23). QuPath provides a user-friendly graphical user interface (GUI), comprehensive instructional documentation and an active and engaged user community, which collectively will help in maintaining and improving DANEELpath. In line with our aim to reach as many users as possible, we have designed this deep learning toolkit to be used via either scripting or an intuitive GUI. Although DANEELpath was initially conceived to extract features from HR-NB GTA-sf models, we demonstrate herein the way these tools are likely to work with other resource material, including different cell cancer types, 3D scaffolds, and human tumor samples. This increases the potential for DANEELpath to be used in other research fields and model systems.

## RESULTS

### The morphometric and staining features of HR-NB cell clusters grown in two GTA-sf scaffoldings exhibit distinctive characteristics

SK-N-BE (2) and SH-SY5Y cell growth in GTA-sf hydrogels resulted in the formation of round cell clusters, a phenomenon we previously observed in PEG and GelMaAlgM-based hydrogels (17,18). The hydrogels exhibited a range of cell cluster sizes and distributions, as observed in both H&E and anti-VN-stained sections. The VN-stained sections revealed disparate patterns in the non-added VN and VN-rich scaffoldings. A distinctive staining pattern was observed in non-VN-added hydrogels, characterized by intra-cluster cytoplasmic staining and frequently pronounced peri-cluster staining, the latter referred to as VN corona (19). In contrast, VN-rich hydrogels exhibited strong matrix staining, resulting from the exogenous addition and appropriate linkage of VN to the GTA-sf. Notably, a faint intensity area was discernible around most clusters in VN-rich hydrogels, which we have called a pale halo (19). Below, we examine the application of DANEELpath tools to facilitate cell cluster segmentation and extraction of spatial and morphometric/staining features to quantify and provide a comprehensive digital analysis of these features.

### A Center-periphery algorithm can achieve equal areas in both locations regardless of ROI shape

Analysis of cell cluster distribution across hydrogel sections revealed a distinct tendency for higher cluster density in the periphery compared to the hydrogel center, particularly evident in the case of SK-N-BE(2) cells. To accurately assess these differences in cluster distribution, we developed an algorithm to precisely delineate the same area of center and periphery ring, as detailed in the Materials and Methods section. This algorithm was applied in hydrogels cultured for 3 weeks with SK-N-BE(2) and SH-SY5Y, as shown in the different examples in Figure 1A. Despite the different shapes, the algorithm found the boundary to obtain a center and periphery of approximately the same area. The absolute percent difference in area between the two regions computed in 48 hydrogel images was less than 0.4%, indicating good accuracy in delineating equal areas (Figure 1B). This algorithm was next used to study the different spatial distribution of cluster densities of each cell line in the presence or absence of VN within the hydrogel, and with or without CLG treatment. In the SK-N-BE (2) cell line, higher cluster density was observed in control (without CLG treatment) than in CLG-treated samples. Moreover, a shift towards the periphery was observed in both control and treated samples, but was more pronounced in VN-rich hydrogels. In contrast, SH-SY5Y cells showed lower cell cluster density with no differences associated with CLG treatment. A slight shift to the periphery of the cell clusters was also observed, although it was less pronounced than in the SK-N-BE (2) cell line.

**Figure 1.**
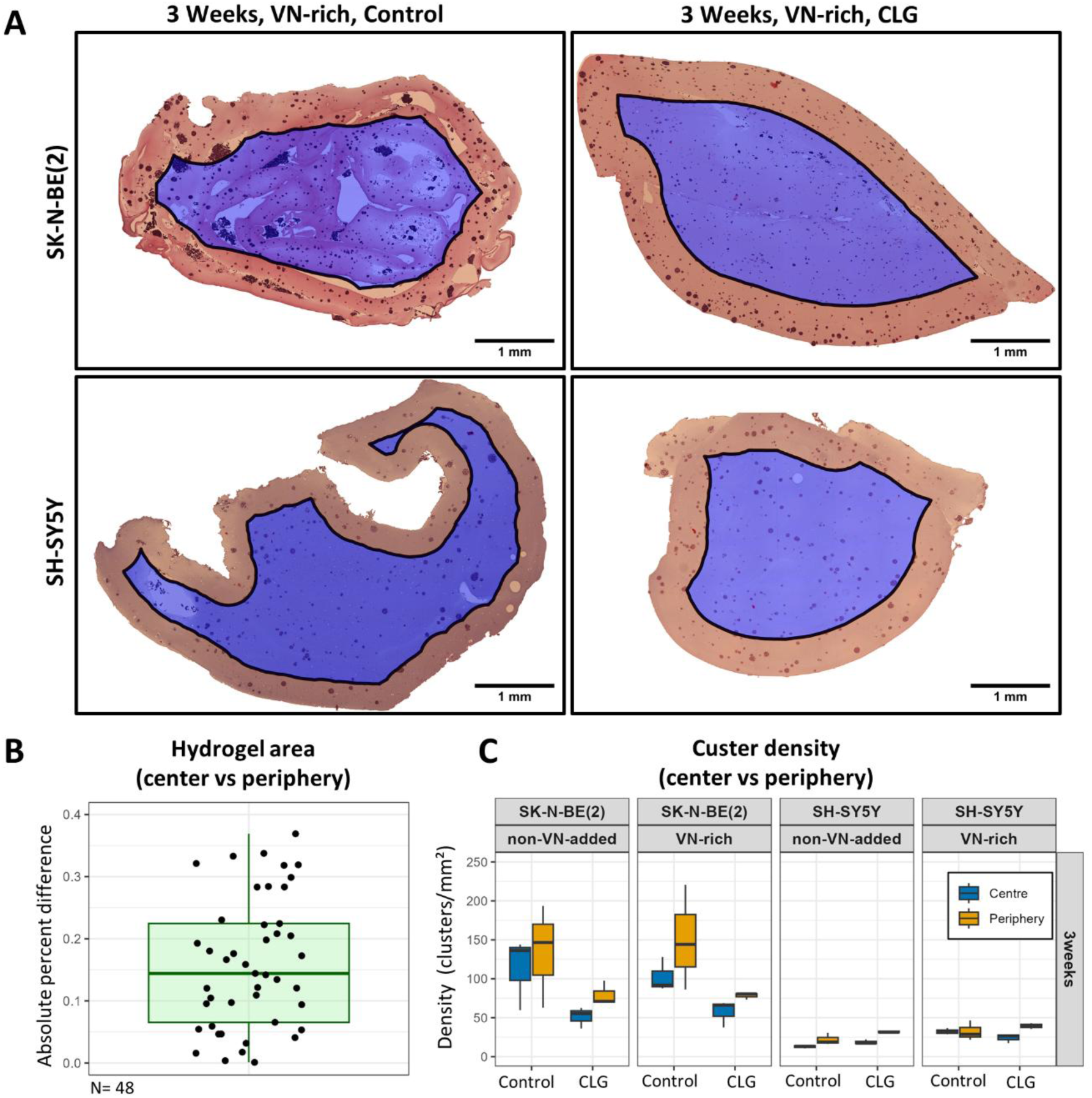
Determination of equal areas of the center and the periphery of the hydrogel. **A**: Hydrogels from the SK-N-BE (2) and SH-SY5Y cell lines in a VN-rich environment without and with CLG treatment after 3 weeks of culture. The algorithm identifies the boundary (black line) separating the center (blue) and the periphery (orange) regardless of hydrogel shape. **B**: Boxplot of the absolute percent difference between the center and the periphery areas in 48 hydrogels. The maximum difference is under 0.4%. **C**: Boxplot of cluster densities at 3 weeks in non-VN-added and VN-rich hydrogels without (control) and with CLG (n=6 hydrogels per condition) reveals differences in the center and periphery between cell lines. Scale Bar: 1 mm. VN: Vitronectin, CLG: Cilengitide.

### Fine-tuning the U-Net training for specific stains enables accurate cluster segmentation

U-Net training for H&E staining was performed for 102 epochs, with micro metrics for IoU of 0.919 and 0.927 and Dice Scores of 0.958 and 0.962 in the validation and test sets, respectively (Figure 2A, i). As expected, the micro metrics for the training set were higher, with an IoU of 0.965 and Dice Score of 0.982 (Figure 2A, i). These values indicate good cluster segmentation with the H&E U-Net, as observed in an example of an H&E-stained patch of the test dataset (Figure 2B).

**Figure 2.**
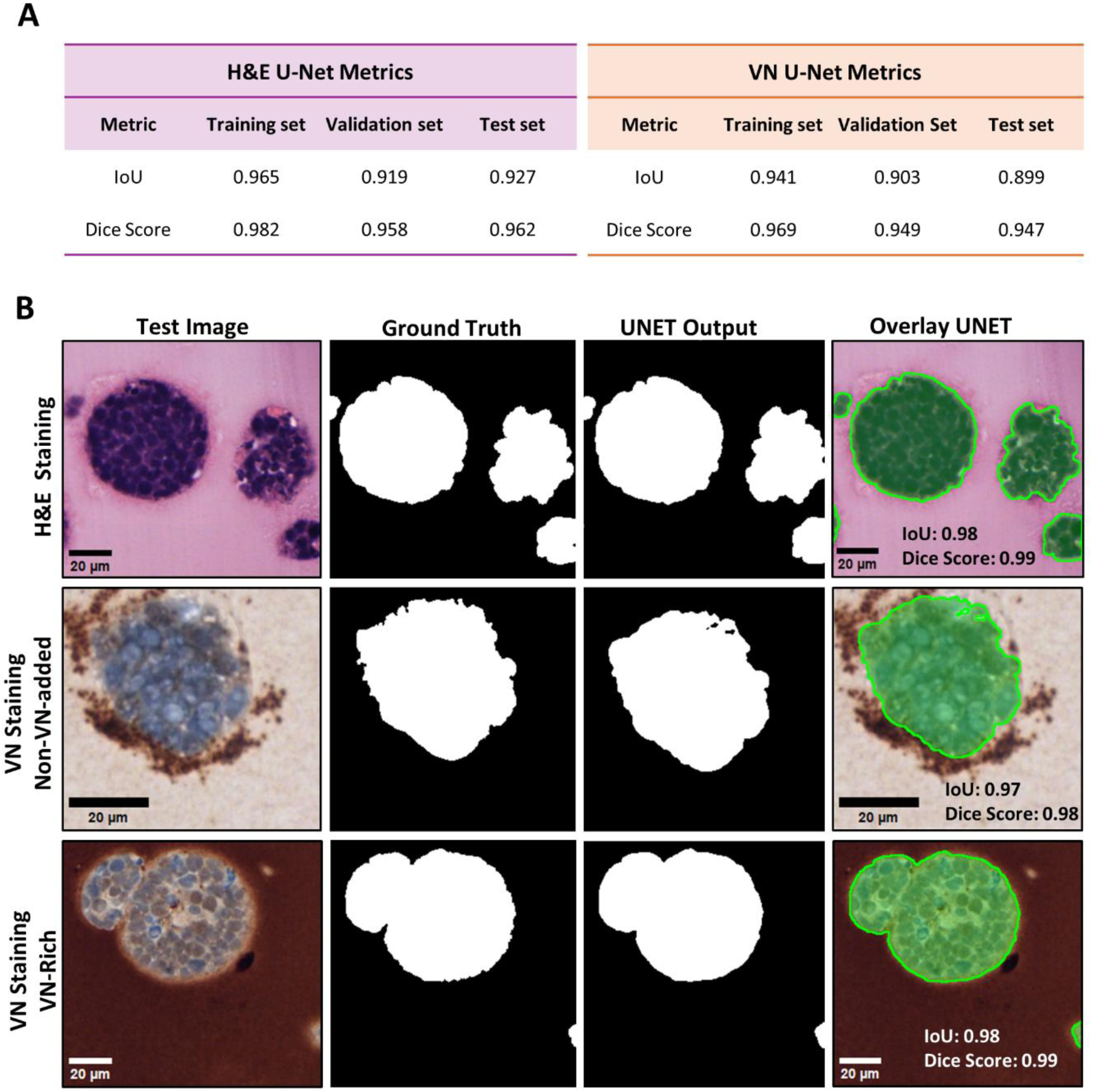
Fine tuning U-Nets for cluster segmentation. **A**: Micro metrics IoU and Dice score metrics computed in the training, validation and test sets for the H&E and VN U-Nets compared with the ground truth. **B**: Examples of semantic segmentation in test images for the H&E and VN U-Nets. Cluster segmentation excludes VN corona and pale halos from VN-rich non-VN-added hydrogels, respectively. H&E: Hematoxylin and eosin, VN: Vitronectin, IoU: Intersection over union.

The VN U-Net model was optimized to circumvent the VN corona and pale halo segmentation, due to the inherent challenges involved in the downstream processing required for associating peri-cluster features with parent clusters. Fine tuning for VN staining U-Net was done after 49 epochs, with micro metric IoU scores of 0.903 and 0.899 and Dice Scores of 0.949 and 0.947 in the validation and test sets, respectively (Figure 2A, ii). As in the H&E U-Net model, the micro metrics for the training set were higher, with an IoU of 0.941 and Dice Score of 0.969. These values suggest a good performance in the cluster segmentation with the VN U-Net, notwithstanding the discrepancy in hydrogel background hue, as observed in test images of non-VN-added and VN-Rich hydrogels stained for anti-VN (Figure 2B).

Overall, the performance of the U-Nets for cluster segmentation was satisfactory for both stains. Some minor issues still required attention, however, such as holes and small matrix miss segmentations outside the clusters. To address this, we implemented a basic post-processing step that included filtering by size, solidity, eccentricity and filling holes (Supplementary Figure 2).

### Average distances between cells and number of nearest neighbor clusters allow detection of potential effects of scaffolds and drug candidates

Cells were successfully detected using the QuPath StarDist extension. The QuPath implementation of DANEELpath tools provides the option to compute the following descriptors for cell distances inside clusters: mean, standard deviation, median, median absolute deviation (MAD), minimum and maximum. Furthermore, DANEELpath has a visualization feature that enables users to draw the connections as detection objects in QuPath. When the mean of the distances between nearest neighbor centroids inside cell clusters were computed in the SK-N-BE(2) cell line [Figure 3A, i], the mean distance to the cluster cell nearest neighbor centroids statistically increased with CLG treatment in both non-VN-added (median ± MAD: control 4.89± 0.35 vs. CLG 5.13 ± 0.43) and VN-rich (median ± MAD: control 4.92 ± 0.41 vs. CLG 5.12±0.46) hydrogels (Figure 3A, ii). This finding suggests that CLG treatment triggers a mechanism that increases the distance between cell centroids in SK-N-BE(2) cells.

**Figure 3.**
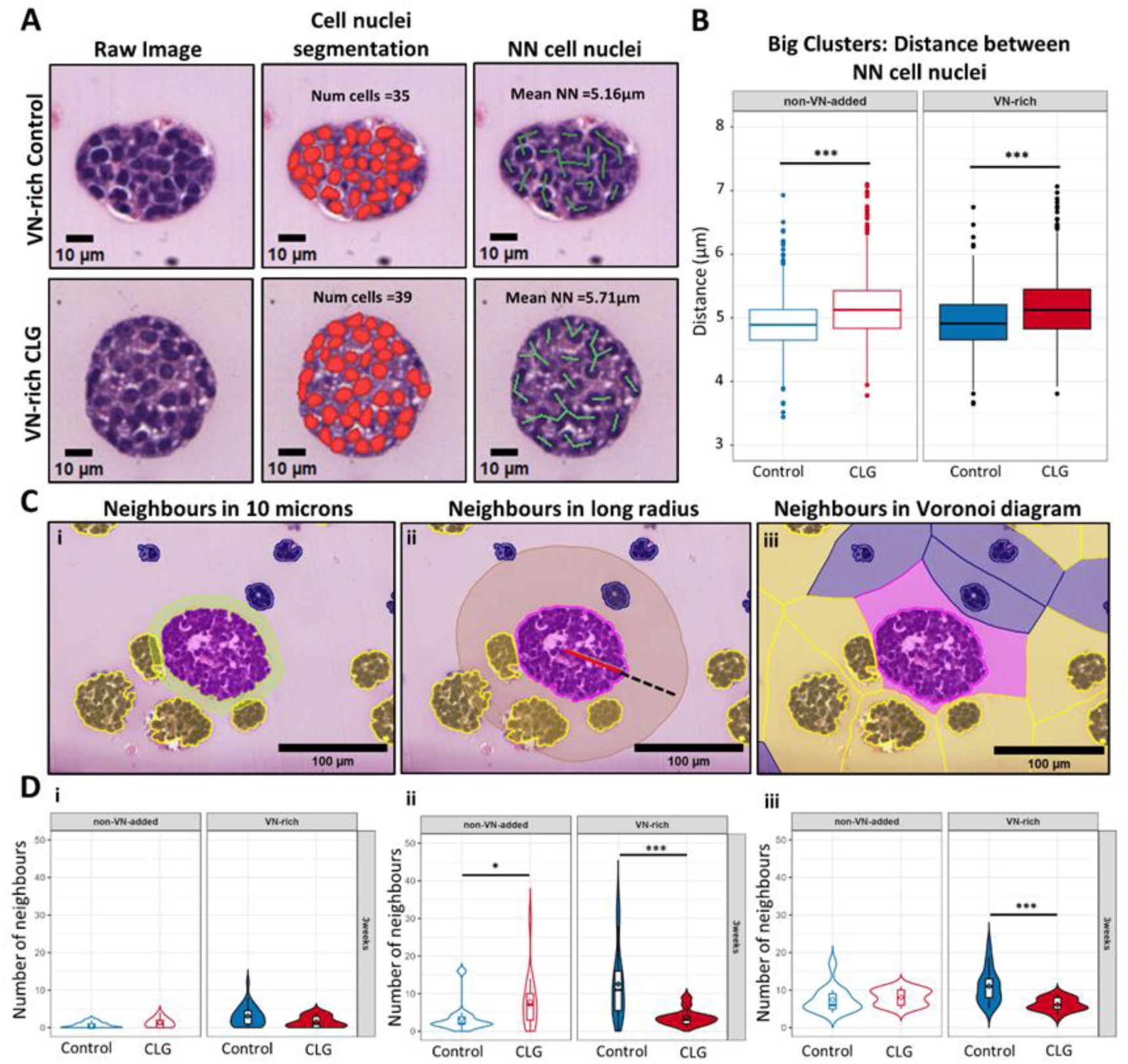
Distance between cells and clusters in hydrogels. **A**: H&E staining of SK-N-BE (2) large clusters in VN-rich hydrogels. Cell nuclei were detected using StarDist. The distance from each cell centroid to its nearest neighbor cell centroid is represented, and the mean of those distances computed. **B**: Boxplot showing the mean distances of cells to the nearest neighboring cell in SK-N-BE (2) big clusters in non-VN-added and VN-rich hydrogels with and without CLG treatment. Statistical analysis using non-parametric Wilcoxon signed-rank test. *** p-value < 0.001. **C**: Cell cluster neighbor counting strategies using a SK-N-BE (2) large cluster (magenta) as reference. **i**: Number of neighbors in 10 microns expansion. **ii**: Number of neighbors in long radius expansion. The red line inside the cluster shows its long radius, while the dashed line shows the expansion. **iii**: Number of neighbors in Voronoi diagram. An influence zone is created from the border of the cluster and the touching expansions are counted. Small and big clusters are marked in blue and yellow, respectively. **D**: Violin plots showing total neighbor counts using three counting strategies for SK-N-BE(2) large cluster cells after 3 weeks of culture in non-added-VN and VN-rich hydrogels with and without CLG treatment. The diamond inside the boxplot shows the mean of the values. Wilcoxon signed-rank test shows differences in cell cluster neighbor counts in long radius and Voronoi methods. *p-value < 0.05, *** p-value < 0.001. VN: Vitronectin, CLG: Cilengitide, NN: Nearest neighbor.

Next, the number of cluster neighbors were computed in the SK-N-BE(2) cell line using three different approaches. For the first approach, an expansion of 10 microns was set (Figure 3B, i) giving a lower number of neighbors. For the long radius (Figure 3B, ii) and Voronoi diagram (Figure 3B, iii) approaches, both yielded the same number of neighbors. A discrepancy in the identity of the neighbors was present. To visualize these differences in neighbor identity, DANEELpath provides a visualization tool that allows users to draw expansions and Voronoi diagrams.

As the next step, the three criteria were compared to obtain the total number of neighbors for large clusters (Figure 3B, iv – vi). For non-VN-added hydrogels, a statistical increase in the total number of neighbors with CLG treatment was observed only with the long radius neighbor method (median ± MAD: control 2 ± 1.48 vs. CLG 7 ±5.93). In contrast, in VN-rich hydrogels, a statistically significant reduction in the total number of neighbors after CLG treatment was detected with the long radius neighbor (median ± MAD: control 11 ± 8.89 vs CLG 3 ±1.48) and Voronoi (median ± MAD: control 11 ± 4.44 vs. CLG 6 ±1.48) methods.

Altogether, the results demonstrate that CLG treatment has a distinct impact on the number of cluster neighbors, in scaffoldings with varying VN amounts. DANEELpath provides users with the flexibility to select the criteria that best fit their biological question.

### The use of Voronoi diagrams and peri-cluster feature segmentation enables the quantification of distinctive staining patterns

Isolating peri-cluster features presents two significant challenges. First, the proximity of two peri-cluster structures makes their isolation difficult, as illustrated in Figure 4A. The second challenge is the association of each peri-cluster structure with the original cluster. A pipeline of analysis was developed to overcome these challenges. We used a Voronoi diagram derived from the cluster border (which creates influence zones) to help resolve the initial challenge, as it effectively delineated the clusters, thereby limiting the peri-cluster structures. Next, an expansion limited by the Voronoi diagram was created to detect the VN corona and pale halo. As the expansion was constructed with JTS, the association of the peri-cluster structure with the original cluster was feasible, thereby addressing the second challenge.

**Figure 4.**
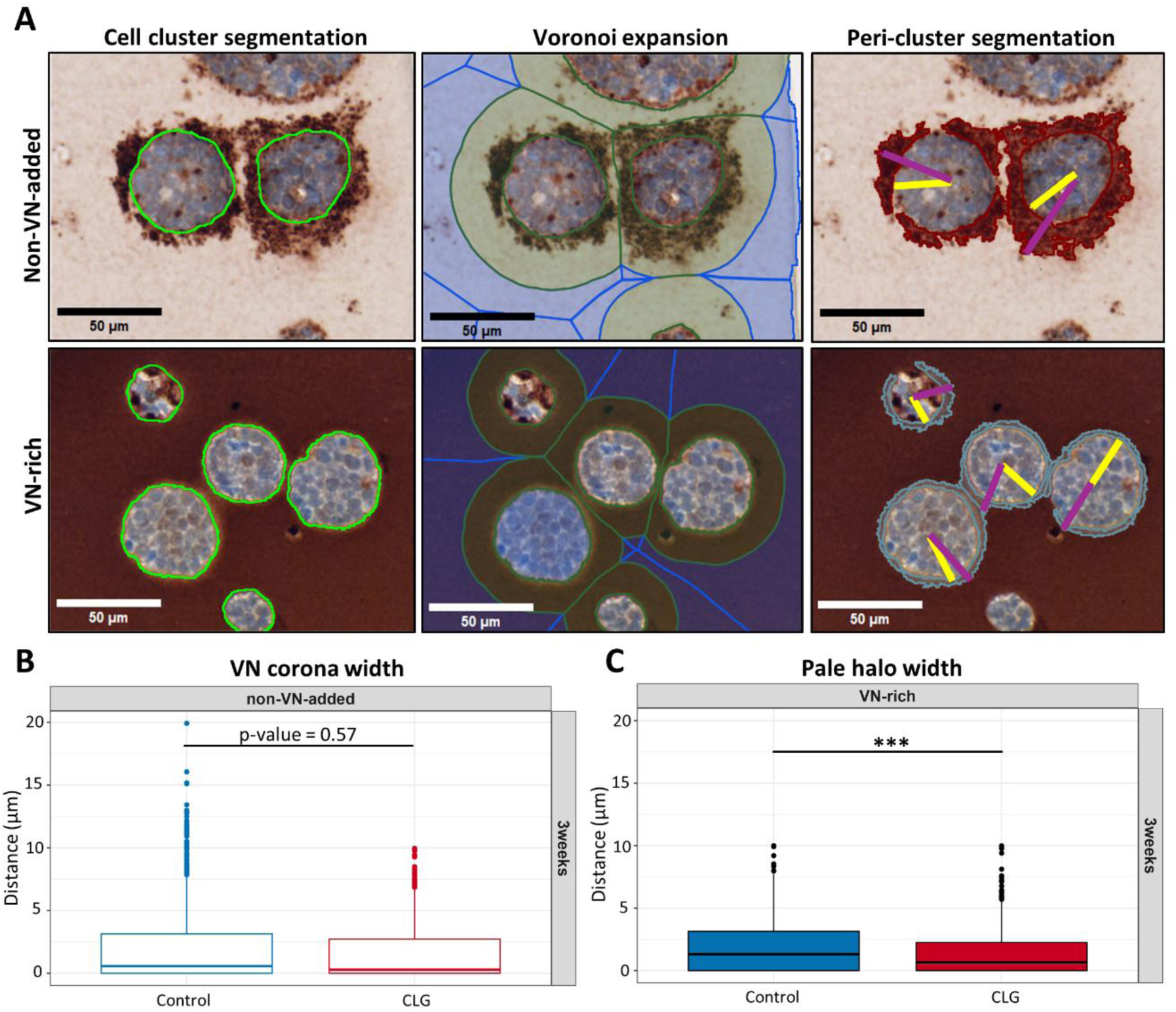
Pipeline for VN corona and pale halo segmentation around clusters. **A**: Once the cluster segmentation has been completed, a Voronoi diagram (shown in blue) is created to define the influence zones of the clusters. A circular expansion from the cluster borders (green expansion) is created and limited to the influence zone. In non-VN-added hydrogels, the VN corona is segmented with a pixel classifier. For VN-rich hydrogels, the pale halo is detected using the triangle auto-threshold method. The width of the VN corona and pale halo is calculated by subtracting the radius of the cluster (shown in yellow) from the radius of the sum of the cluster and peri-cluster structure (shown in magenta). **B**: Boxplot representing the width of VN corona in SK-N-BE (2) big cluster cells at 3 weeks of culture in non-VN-added hydrogels. A tendency towards reduced VN corona width is observed in CLG-treated hydrogels. **C**: Boxplot representing the width of the pale halo in SK-N-BE (2) big cluster cells at 3 weeks in VN-rich hydrogels. A decrease in pale halo width is observed in CLG-treated hydrogels. Statistical analysis using non-parametric Wilcoxon signed-rank test. *** p-value < 0.001. VN: Vitronectin, CLG: Cilengitide.

We then computed the VN corona width for big clusters in non-VN-added SK-N-BE (2) hydrogels (Figure 4B). A trend towards decreased VN corona width was observed after CLG treatment (median ± MAD: control 0.57 ± 0.85 vs 0.28 ± 0.42). In VN-rich hydrogels, the width of the pale halo was quantified. Interestingly, we observed a statistically significant decrease in the pale halo from CLG-treated hydrogels (median ± MAD: control 1.31 ± 1.94 vs CLG 0.66 ± 0.98). Overall, a reduction in both the VN corona and the pale halo was observed after CLG treatment.

### DANEELpath tools are applicable to a wide range of other scenarios

DANEELpath tools were developed specifically to facilitate analysis of HR-NB cell GTA-sf hydrogels; however, their applicability to other samples has also been explored, as illustrated in Figure 5. Figure 5A provides an example of a group of cells from a HR-NB immunostained sample against the macrophage marker CD68, which revealed a tendency for CD68-positive cells to be located towards the periphery of the tumoral niche. Therefore, the DANEELpath algorithm was used to define the central and peripheral regions, automatically calculating number of CD68-positive cells in both locations of the tumoral niche. Another potential application of DANEELpath is the use of the H&E U-Net in other hydrogel models (Figure 5B). In GTA-sf Ewing sarcoma models, the H&E U-Net appears to function effectively for cluster segmentation. The model was further tested on images from a previous study, using GelMA-ALG 3D-printed models. Despite the changes in the hydrogel background color resulting from the 3D printing process, the U-Net model performed acceptably in the segmentation task. The VN U-Net was also tested in HR-NB cell GTA-sf hydrogels stained against the proliferation nuclear cell marker Ki67 and the cytoplasmic antiapoptotic marker Bcl2 (Figure 5C), with excellent results. Tissue microarray (TMA) cores of HR-NB human samples and orthotopic mouse tumors were also utilized and analyzed with DANEELpath tools. In this instance, the cores were stained against the CD31 marker, which highlighted the blood vessels. Subsequently, the Voronoi diagrams feature in DANEELpath was employed to create the influence zones of the segmented vessels. The vessels and the Voronoi diagram were transferred to an image of the same TMA core but stained against VN. Once this data is aligned, we can validate the previously described topological features of the VN related to blood vessels (36) according to the vessel’s influence zones as defined by DANEELpath. Collectively, these examples demonstrate the applicability of DANEELpath to the detailed analysis of morphometric and staining features in models other than GTA-sf hydrogels of HR-NB cells.

**Figure 5.**
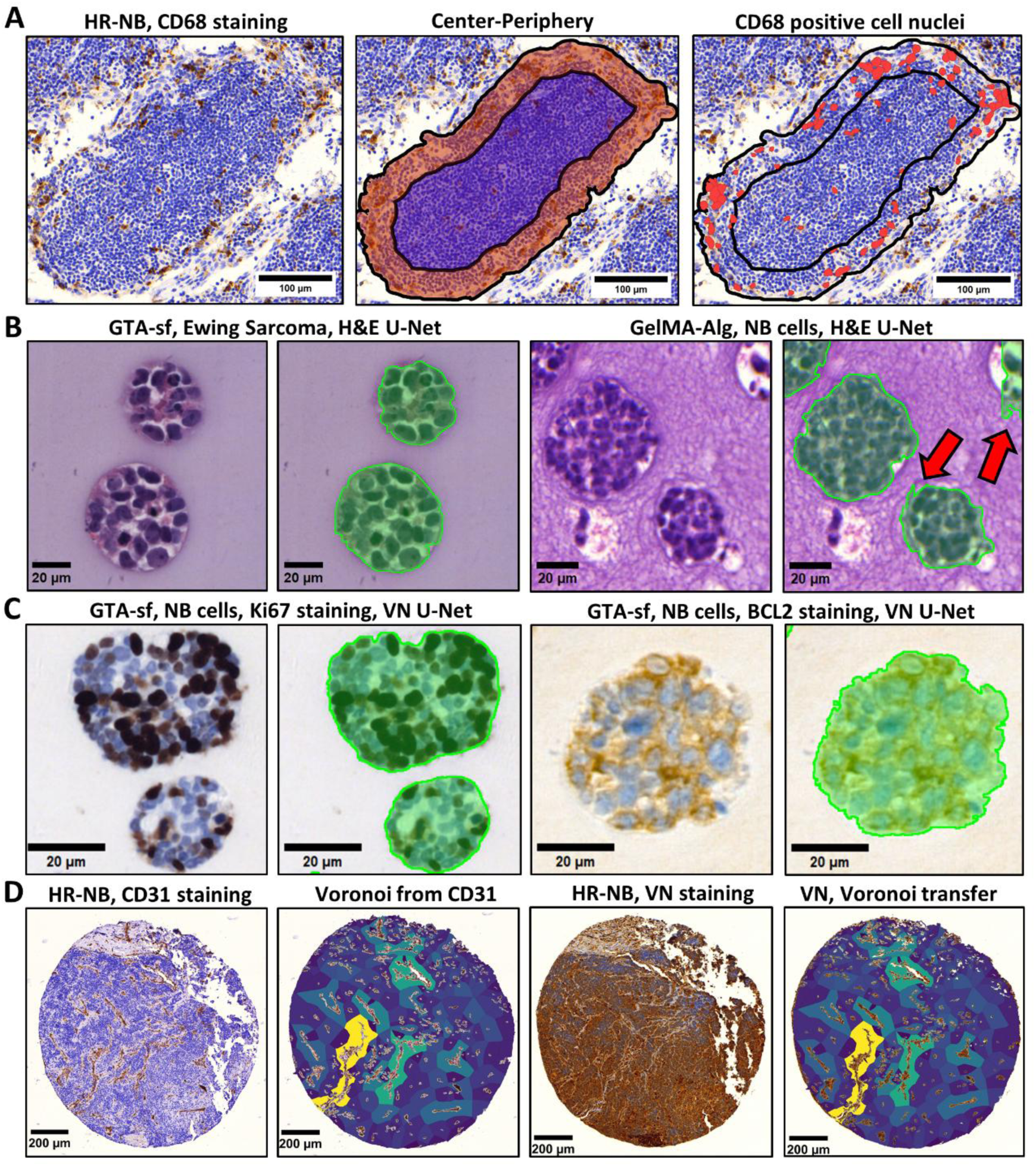
Other scenarios for utilizing DANEELpath. **A**: Assessment of the center (blue) and the periphery (orange) in a group of cells from a human HR-NB sample immuno-stained with CD68 (red, macrophage). CD68-positive cells are located mainly at the periphery. **B**: Cluster segmentation using H&E U-Net in Ewing Sarcoma cell lines grown in GTA-sf hydrogels. **C**: Cluster segmentation using H&E U-Net in SK-N-BE (2) cells grown in GelMA-Alg hydrogels. H&E U-Net can segment NB clusters in hydrogels of different composition with minor errors (red arrows). **D**: SK-N-BE (2) cells in GTA-sf hydrogels immuno-stained against Ki67 and BCL2. VN U-Net is able to segment the clusters with nuclear and cytoplasmic staining. **E**: Correlation between blood vessels and VN expression in HR-NB samples. From a HR-NB core stained against CD31 a Voronoi diagram is obtained to represent the blood vessel area and create influence zones. The influence zone color indicates the number of touching neighbors. Next, a consecutive section core of the HR-NB sample is immuno-stained against VN to transfer the CD31 influence zones to the VN stained core using affine registration. HR-NB: High-risk neuroblastoma, H&E: Hematoxylin and eosin, GTA-sf: Gelatin-tyramine with silk fibroin, GelMA-Alg: Methacrylated gelatin/alginate, VN: Vitronectin.

## DISCUSSION

While 3D hydrogels are an effective tool for studying the role of ECM elements in HR-NB, analyzing 3D hydrogel images, particularly cell cluster segmentation, is a significant challenge that hinders the ability to conduct high-throughput analysis with a greater number of candidate drugs and cell lines. By developing the DANEELpath toolkit we could streamline these samples and provide tools to explore morphometric and staining features at the level of cells and clusters, using WSI.

In pathology, the proportion of central and peripheral regions in a delimited lesion or tumor is difficult to define with a microscope. DANEELpath enables an automatic determination of these two regions independently of the original shape, and in our research this has proven to be robust, showing a maximum difference between regions of 0.4% in 48 WSI hydrogel samples. Furthermore, DANEELpath was capable of handling multiple ROIs within a single image, thereby avoiding the need to isolate them as individual images, unlike in our previous implementation (27). The concept of center and periphery are readily comprehensible and may serve as a means of standardizing spatial definitions in both hydrogels and human samples. In the present study, this approach was demonstrated in human samples, where a higher distribution of CD68-positive cells was observed at the periphery of a group of neuroblastic cells within an HR-NB sample. This illustrates that defining these locations can help formulate biological questions regarding the underlying cause of this distribution. Nonetheless, numerous additional applications are possible, such as defining center-periphery regions to investigate disparities in hypoxic regions within hydrogels and human samples by guiding laser micro-dissections and performing RNAseq (37).

Our fine-tuned U-Net, supported by Efficient-NetB4, demonstrated effective cluster segmentation in both H&E and VN-stained hydrogel samples, irrespective of VN concentration and the wide range of cell cluster size. For matrices with varying compositions, such as GelMA-Alg, segmentation was successful, although there were some errors in cluster border delineation. The Efficient-Net family has performed excellently in other medical fields such as breast cancer (38), skin cancer (39,40) and lung and pneumothorax segmentation (41,42). Our future work plan also includes creating a large training dataset comprising additional staining, cell lines, and images from Gel-MA and PEG models and designing a pipeline to streamline 3D reconstruction from these 2D images. The flexibility of DANEELpath allows easy adaptation of inference for multiclass semantic segmentation. In this context, we are currently adapting DANEELpath U-Net architecture to distinguish tumor parenchyma from stroma in high-resolution human neuroblastoma samples. The ultimate goal is to make the DANEELpath implementation suitable for use in 3D model experiments and human sample analysis.

In terms of the number of neighbors in clusters, we have outlined three approaches. Our analysis revealed that CLG exerts a different effect on spatial distribution according to whether VN is incorporated into hydrogels. This reflects the importance of ECM composition and the influence of mechano-drugs on cluster growth dynamics. Neighborhood analysis in tissue has a wide range of applications in biomedical research. It has been used to assess rearrangements in ECM in colorectal cancer (43), to define the normal liver lobular architecture (44), and in modeling wound healing processes (45).

While DANEELpath functionality was originally designed to assess cluster distribution, we also explored its potential use in human samples. We demonstrated that the Voronoi diagram can be created from blood vessels and can be used to investigate the topological characteristics of VN within blood vessel influence zones, as previously reported (36). The perivascular microenvironment represents a promising area of interest, particularly in light of recent studies that have identified potential therapeutic targets within this niche. These include perivascular macrophages, as well as the perivascular niche itself, which encompasses tumor-associated endothelial cells, mural cells and cancer-associated fibroblast (46,47). Delineating the influence zones associated with these elements may bring valuable insights into the perivascular TME of NB. We demonstrated that applying Voronoi diagrams facilitated the acquisition of morphometric features of peri-cluster structures. Furthermore, this application of DANEELpath may prove valuable in addressing other biological inquiries. For example, it can be used in osteoarthritis studies to evaluate alterations in the peri-cellular matrix and metalloproteinases secretion surrounding chondrocytes (48).

We developed DANEELpath using tools already built within QuPath, as it is currently one of the most widely used open-source software options, offering intuitive GUI, comprehensive documentation, and robust support for working with WSI. Despite incorporating deep learning, therefore, no further installations are required for other functionalities. U-Net requires creating an Anaconda environment, primarily in order to streamline U-Net deployment and provide the flexibility to deploy new state-of-the-art models in the near future. The use of these environments further allows users to benefit from CUDA GPU acceleration when available. Finally, two versions of the functionalities were developed, accompanied by installation instructions accessible via GitHub repository (https://github.com/iviecomarti/DANEELpath). The script version allows users with medium level experience to utilize the tools with greater flexibility, for multiple images simultaneously. The GUI version allows users without programming experience to make use of DANEELpath tools within the existing QuPath image. In conclusion, the creation of user-friendly and comprehensive DANEELpath image analysis tools may facilitate their uptake within the scientific community for implementation in *in vitro* and *in vivo* models, and human samples, thereby helping to advance biomedical discovery.

## MATERIALS AND METHODS

### Experimental models

To develop the methods, we used an in-house dataset comprising HR-NB cell 3D hydrogels. Briefly, from a variety of cell lines available, we chose *MYCN*-amplified SK-N-BE(2) and *ALK*-mutated SH-SY5Y human NB cell lines, since these two tumor types represent 64% (50 and 14%, respectively) of HR-NB (24,25). NB cell lines were acquired from American Type Culture Collection (ATCC, Manassas, VA, USA) and cultured for 2 and 3 weeks in GTA-sf hydrogels following Hasturk O *et al.*’s methodology (26), with and without adding VN to the scaffolding and with vs. without CLG treatment (three or six doses of 100 µM, respectively) as employed previously in 2D cell lines (21). SK-N-BE(2) and SH-SY5Y cell lines secreted VN to culture media when grown in 2D and 3D models (16). The data set yielded four groups: hydrogels without added VN (non-VN-added), containing NB cell-secreted VN only, hydrogels with added VN (VN-rich), hydrogels without added VN and with CLG (non-VN-added-CLG) and hydrogels with added VN and CLG (VN-rich CLG). The samples were FFPE and 3-micron slides were stained with Hematoxylin & Eosin (H&E) and immunostained against VN (anti-VN) [clone EP873Y, isotype IgG, code ab45139, Abcam].

A total of 96 WSI (48 slides per cell line/stain) were digitized with the Roche Ventana iScan HT 20x (32 H&E stains and 31 anti-VN stains) and the 3D Histech PANNORAMIC 250 Flash III 40x (12 H&E stains and 17 anti-VN stains). Cell clusters were annotated using pixel classifiers, and when necessary were manually adjusted on QuPath. A total of 26392 cell clusters were segmented for H&E and 23267 for anti-VN staining, including their coronas and surrounding pale halo. Segmentations were validated by a histologist (RN) and a pathologist (SN).

### Determination of hydrogel center and periphery

To study the size distribution of cell clusters, the same area of center and periphery ring was defined, as previously reported (27). To achieve this in any two-dimensional (2D) closed shape, we implemented a slight improvement from our previous work. First, we obtain the area (pixels^2^) and the perimeter (pixels) of a Region of Interest (ROI), namely the hydrogel. Next, the distance that must be reduced from the borders of the ROI to achieve a dilation factor **“n”** of ½ is calculated as **’x’** using **Formula 1:**

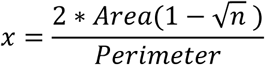

The mathematical logic behind **Formula 1** can be found in **Supplementary Figure S1**. The initial expansion can be created with Java Topology Suite (JTS) tools in QuPath. If the absolute percent difference between the initial expansion and the target area (**ROI area/2**) is higher than 1%, a while loop minimizes this difference. Finally, subdivisions are created and added as annotation objects in QuPath.

### Fine-tuning U-Net training for cell cluster segmentation

Cell clusters are the units of interest for the extraction of morphological and spatial information. One of the main challenges is the significant variation in cluster size, ranging from single cell clusters to those containing from 1 cell to 926 cells. This wide size distribution makes it difficult to use tools such as StarDist, Cellpose, or InstanSeg, as these methods are primarily designed to segment cell instances rather than cell clusters. Therefore, to streamline the process of cell cluster segmentation for cultures, two U-Nets for semantic segmentation were fine-tuned: one for H&E and one for anti-VN staining. The images and labeling were divided into 256×256 pixel tiles. For data augmentation purposes, image resolution was not standardized. The H&E staining model was trained on a dataset comprising 21664 tiles, with 2708 reserved for validation and test sets. The VN staining model was trained on a dataset comprising 26995 tiles, with 3374 reserved for the validation and test sets.

We selected a U-Net architecture (28) with an EfficientNet-B4 encoder (29) pre-trained on the ImageNet dataset (30), as provided by the Segmentation-Models-PyTorch package (31). The PyTorch framework (pytorch.org) was employed in conjunction with the PyTorch-Lightning package (lightning.ai) to finetune the model. During the training process, on-the-fly data augmentation was applied using the Albumentations package (albumentations.ai), which included vertical and horizontal flips, adjustments for brightness and contrast, hue, saturation, and Contrast Limited Adaptive Histogram Equalization (CLAHE).

In accordance with previously established criteria (32), models were evaluated using the micro-metrics of the Dice score and the Intersection Over Union (IoU). The Dice score and IoU were calculated using the values of true positives (TP), false positives (FP), and false negatives (FN), as follows:

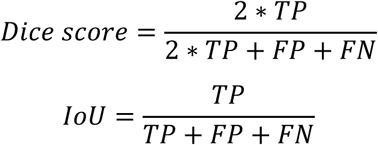

Models were trained for 200 epochs with a batch size of 32. The Adam optimizer was selected, starting with a learning rate of 0.0002, which was gradually reduced to 0.00001 via a Cosine Annealing Scheduler (arXiv:1608.03983). The validation loss was monitored, and early stopping was applied if no improvement was observed for 20 epochs.

### Distance between cell centroids inside clusters

To evaluate the impact of VN or CLG on cell arrangement within clusters, the distance was measured from each cell centroid to its nearest neighboring cell centroid within the cluster. Cells were detected using the StarDist extension in QuPath (4). Next, each cell’s nearest neighbor was identified and the Euclidean distance between their centroids was calculated using QuPath Delaunay tools. To summarize the results, the mean distances for cells within each cluster were computed.

### Cell cluster neighbor counting

Cell clusters were classified into three groups based on the number of cells present: small (1–10 cells), medium (11–95 cells), or large (95–926 cells). To analyze the spatial relationships between these clusters, the number and classification of neighboring cell clusters were also counted.

Three methods were implemented to define the neighborhood of a cell cluster using JTS. In the first method users can specify a fixed distance in microns and count the number and classification of neighboring cell clusters within this specified distance. With the second method, the longest radius of each cluster is expanded and the number of clusters and their classification within this radius are counted. For those two methods, any cluster located within or touching this expanded area is counted as a neighbor. The third method involves creating Voronoi diagrams from the borders of the cell clusters (also known as influence zones [33]). This method counts the touching neighbors along with their classifications according to the Voronoi annotations, similar to the pyclesperanto python package (34).

### VN corona and pale halo measurements

The width of VN corona secretion (in non-VN-added) and pale halos (in VN-rich scaffolds) in WSI anti-VN stains were evaluated relative to the cell cluster. To achieve this, Voronoi diagrams were created from the borders of the cell clusters. Subsequently, a concentric expansion was conducted around the cell cluster, with the boundaries constrained by the Voronoi diagram. In this way, cell cluster expansion does not encroach upon the influence zone of other clusters. To perform the VN corona segmentation, a simple thresholding method was employed, utilizing the average channel within the cluster expansion. Pale halo segmentation was performed using the triangle auto-threshold for each expansion with a custom Groovy script. The VN corona or pale halo was then merged with the original cluster using JTS, and the radius calculated for the merged structure. To obtain the width of the VN corona and the pale halo, the radius of the cluster was subtracted from the merged structure.

### QuPath implementation of developed methods

U-Nets were run from QuPath using the Virtual Environment Runner developed by BIOP (35). Once the inference was executed in QuPath, the tiles were exported, and a command sent to the terminal to run a Python inference script. When the inference was complete, the segmentations were imported back to QuPath, where post-processing could be applied. The advantage of this method is that it facilitates the use of graphics processing unit (GPU) acceleration. U-Net architecture fits in an entry level NVIDIA GPU with 4GB of VRAM, avoiding the need for heavy investment. In QuPath, the user can set the batch size to decrease the number of loadings between the central processing unit (CPU) and GPU. If GPU acceleration is not available, the inference script automatically uses the CPU for inference. To facilitate the use of as many developed tools as possible with QuPath, a graphical user interface version has been created for users without coding experience (including U-Net). Furthermore, the Groovy script version for each tool has also been provided, enabling multiple images to be processed. The code is available on GitHub, accompanied by instructional videos.

### Statistics Analysis

Data were analyzed and represented using R and RStudio (4.2.2, R Foundation for Statistical Computing, Vienna, Austria). To test the difference in the distance between cell centroids, the difference between the neighborhood counts and the width of VN corona and pale halo, a non-parametric two-tailed Wilcoxon signed-rank test was used. The differences were considered significant with a p-value < 0.05.

### Ethics statement

This study focuses in the development and application of digital pathology pipelines for neuroblastoma 3D models. At the end of the article we show how the algorithms may be used in human samples images. Our laboratory is Spanish Reference Centre for NB Molecular and Pathological studies (Department of Pathology, University of Valencia-INCLIVA) and we have stored in our biobank (reference B.0000339 29/01/2015) samples, imaging data and genetics data. This study was approved by the Research Ethics Committee of the Clinic Hospital of Valencia (No. 2020/025, Act: 372, 30 September 2021).

## ACKNOWLEDGEMENTS

The authors gratefully acknowledge funding support from ISCIII (FIS) and FEDER (European Regional Development Fund), grant number PI20/01107, FNB (Fundación Neuroblastoma), grant number PRV/00166 and CIBERONC (contractCB16/12/00484). IVM was supported by the Ministry of Science, Innovation and Universities of Spain (FPU20/05344). The authors thank Kathryn Davies for English correction. Finally, we appreciate the help of the QuPath Team and the Image.sc forum community in resolving doubts during the development of DANEELpath.

## AUTHOR CONTRIBUTIONS

**Conceptualization:** I.V.M and R.N.

**Investigation:** I.V.M, S.G.A

**Software:** I.V.M

**Methodology:** I.V.M, S.G.A., A.L.C, S.N and R.N.

**Writing – original draft:** I.V.M, S.G.A. and R.N.

**Writing – review and editing:** All authors have read and approved the final version of the manuscript.

