## Supplemental Figures for "DANEELpath: Open-source Digital analysis tools for histopathological research. Applications in NEuroblastoma modELs"

**A**

**i)** Scale functions of area and perimeter for a 2D shape. Change constant **c** to $\lambda s$:

$$A(c) = c^2A_0 \qquad\qquad A(s) = \lambda^2 s^2 A_0$$
$$P(c) = cP_0 \quad \longrightarrow \quad P(s) = \lambda s P_0$$

**ii)** A'(s) == P(s):

When $\lambda = \frac{P_0}{2A_0}$ $\longrightarrow$ $A'(s) = 2s\lambda^2 A_0 = \lambda s P_0$= P(s)

**iii)** Obtention of value of **s**:

$$A_0 = s^2\lambda^2 A_0 \longrightarrow 1 = s^2\left(\frac{P_0}{2A_0}\right)^2 \longrightarrow s = \frac{2A_0}{P_0}$$

**iv)** Decrease of **s** to obtain the **n**th scaling factor:

$$nA_0 = (s - x)^2\lambda^2 A_0 \longrightarrow n = \left(\frac{2A_0}{P_0} - x\right)^2 \left(\frac{P_0}{2A_0}\right)^2$$

$$\boxed{x = \frac{2A_0(1 - \sqrt{n})}{P_0}}$$

**B**

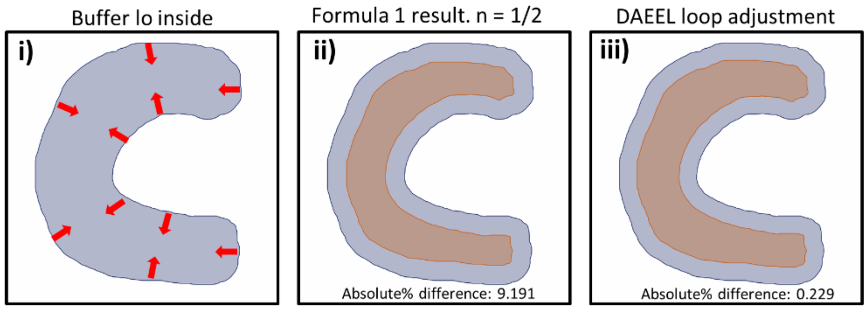


**Figure S1:** Formula 1 logic and DANEELpath implementation. **A**: Mathematical explanation of Formula 1. **I**: First, functions to scale initial area ($A_0$) and perimeter ($P_0$) for a 2D shape with respect to the constant **c**. Then constant **c** is changed to constants $\lambda s$. **ii**: Change of variable makes A'(s) == P(s) possible for a value of $\lambda$. **iii**: the value of constant **s** is obtained by changing the value of $\lambda$ in the case $A_0$. **iv**: Determination of how much distance **x** from the border of the region of interest is needed to decrease constant **s** to obtain a scaling factor of **n**. The valid solution of **x** for this purpose is Formula 1. **B**: DANEELpath implementation. **I**: DANEELpath buffers to inside the original 2D shape (blue) as represented by red arrows. **ii**: First buffering of Formula 1 for n= ½. The absolute percent difference between the center (orange) and periphery (blue) is 9.191. **iii**: DANEELpath loop adjusts the expansion to minimize the difference. Final absolute percent difference is 0.229.

**A**

| H&E U-Net metrics summary per tile | | |
|---|---|---|
| Metric | Validation set (mean ± SD) | Test set (mean ± SD) |
| IoU | 0.863 ± 0.215 | 0.867 ± 0.214 |
| Dice Score | 0.905 ± 0.194 | 0.907 ± 0.193 |

| VN U-Net metrics summary per tile | | |
|---|---|---|
| Metric | Validation set (mean ± SD) | Test set (mean ± SD) |
| IoU | 0.832 ± 0.221 | 0.823 ± 0.228 |
| Dice Score | 0.884 ± 0.205 | 0.877 ± 0.212 |

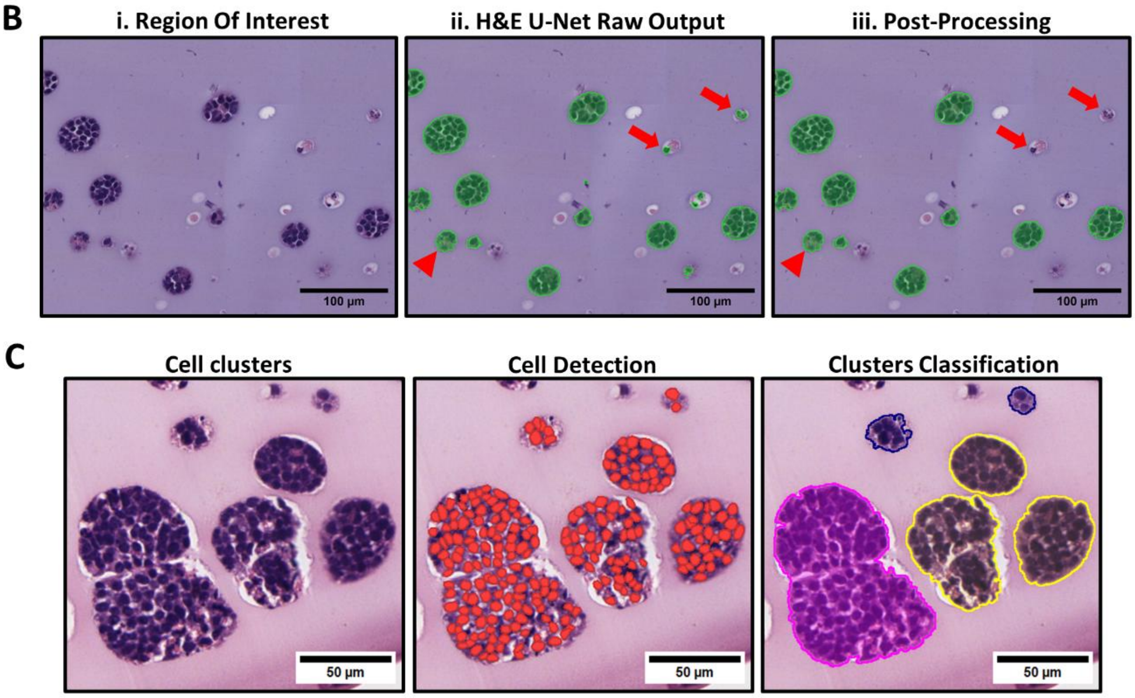


**Figure S2:** Additional information from trained U-Nets. **A**: Summary of the IoU and Dice Score per tile metrics in the validation and test datasets for H&E and VN U-Nets. **B**: Example of inference with post-processing. **i**: Region of Interest. **ii**: Segmentation of H&E model with a tile padding of 96. DANEEL handles 256x256 tiles trimming with the padding and imports the results to QuPath. Cluster mis-segmentations (arrows) and small holes (arrowhead) are observed. **iii**: Post-processing including “fill holes", "minimum area size of 100µm2 “, “maximum eccentricity of 0.9” and “minimum solidity of 0.3”. **C**: Cell cluster classification according to the number of cells. Cell nuclei were detected using StarDist. Clusters are classified as small (blue, 1–10 cells), big (yellow, 11–96 cells), and giant (magenta 96–926 cells).
